# RNA-DNA hybrids support recombination-based telomere maintenance in fission yeast

**DOI:** 10.1101/458968

**Authors:** Yan Hu, Henrietta W. Bennett, Na Liu, Martin Moravec, Jessica F. Williams, Claus M. Azzalin, Megan C. King

## Abstract

A subset of cancers rely on telomerase-independent mechanisms to maintain their chromosome ends. The predominant “alternative lengthening of telomeres” pathway appears dependent on homology-directed repair (HDR) to maintain telomeric DNA. However, the molecular changes needed for cells to productively engage in telomeric HDR are poorly understood. To gain new insights into this transition, we monitored the state of telomeres during serial culture of fission yeast (*Schizosaccharomyces pombe*) lacking the telomerase recruitment factor Ccq1. Rad52 is loaded onto critically short telomeres shortly after germination despite continued telomere erosion, suggesting that recruitment of recombination factors is not sufficient to maintain telomeres in the absence of telomerase function. Instead, survivor formation coincides with the derepression of telomeric repeat-containing RNA (TERRA). In this context, degradation of TERRA associated with the telomere in the form of R-loops drives a severe growth crisis, ultimately leading to a novel type of survivor with linear chromosomes and altered cytological telomere characteristics, including the loss of the shelterin component Rap1 (but not the TRF1/TRF2 orthologue, Taz1) from the telomere. We demonstrate that deletion of Rap1 is protective in this context, preventing the growth crisis that is otherwise triggered by degradation of telomeric R-loops in survivors with linear chromosomes. These findings suggest that up-regulation of telomere-engaged TERRA or altered recruitment of shelterin components can support telomerase-independent telomere maintenance.

## INTRODUCTION

Telomeres are specialized nucleoprotein structures at the ends of linear eukaryotic chromosomes. Telomeres protect the chromosome ends from degradation, aberrant recombination and end-to-end fusions {de Lange, 2009 #4;Dehe, 2010 #444}, while also supporting the formation of subtelomeric heterochromatin {Buhler, 2009 #424}. Telomeric DNA consists of tandem double-stranded G-rich repeats and a singlestranded 3’ overhang {Blackburn, 1991 #425}. The 3’ ssDNA overhang is sequestered by engaging in higher-order structures in part due to the binding of sequence-specific telomeric binding proteins {Blackburn, 2001 #426}. In at least some organisms, the telomere binding protein TRF2 drives formation of a structure called “t-loop”; TRF2 is a component of the shelterin complex that includes the additional factors TRF1, RAP1, TIN2, TPP1 and POT1 in mammalian cells {de Lange, 2005 #17}. The fission yeast (*S. pombe*) genome encodes several classes of telomere binding proteins, many of which are orthologous to mammalian shelterin components. For example, Taz1 (related to mammalian TRF1/TRF2) recognizes dsDNA telomere repeats {Cooper, 1997 #40}, while the conserved Pot1 specifically recognizes the 3’ ssDNA G-rich overhang {Baumann, 2001 #41}. These modules are bridged by the shelterin components Rap1 {Chikashige, 2001 #403;Kanoh, 2001 #404}, Poz1 and Tpz1 {Miyoshi, 2008 #42}; Rap1 constitutes the binding interface with Taz1 {Chikashige, 2001 #403;Kanoh, 2001 #404}. Cells lacking Taz1 undergo telomere lengthening {Cooper, 1997 #40}, due at least in part to increased telomerase recruitment {Dehe, 2012 #402;Moser, 2011 #360}. In contrast, loss of Pot1 drives rapid telomere loss {Baumann, 2001 #41}, likely due to extensive 5’ end resection {Pitt, 2010 #45}.

In fission yeast, telomeric DNA is composed of ~300 bps of degenerate GG_1-5_TTAQA] repeats {Cooper, 2006 #428}. Telomere length is influenced by numerous factors, including mechanisms regulating telomere replication {Maestroni, 2017 #419;Martinez, 2015 #418}, telomerase recruitment and activation {Armstrong, 2017 #420;Nandakumar, 2013 #10}, and the protection of the chromosome end from nucleases {Longhese, 2010 #430;Shore, 2009 #431}. The primary mechanism to counteract telomere loss is the action of the reverse transcriptase telomerase. The fission yeast telomerase is comprised of the conserved catalytic subunit Trt1 {Nakamura, 1997 #19}, the RNA component TER1 that encodes the template for addition of the degenerate telomeric repeat to support telomere elongation {Leonardi, 2008 #22;Webb, 2008 #23}, the conserved subunit Est1 {Beernink, 2003 #21}, and accessory factors such as the Lsm proteins {Tang, 2012 #371} and the La-like protein LARP7/Pof8 {Collopy, 2018 #416;Mennie, 2018 #415;Paez-Moscoso, 2018 #414}. In fission yeast, the factor Ccq1 links telomerase recruitment to the 3’ telomeric overhang through Tpz1 and Pot1 to support telomere maintenance {Tomita, 2008 #1;Miyoshi, 2008 #42}.

Somatic mammalian cells do not display telomerase activity at sustained levels {Kim, 1994 #432;Wright, 1996 #433}, primarily due to transcriptional silencing of the catalytic subunit TERT {Meyerson, 1997 #417;Nakamura, 1997 #19}, and therefore can undergo a limited number of cell divisions before critically short telomeres drive cell cycle arrest and senescence or apoptosis. Thus, cancer cells typically harbor mutations that lead to a gain of telomerase activity {Kim, 1994 #432;Shay, 2012 #47} likely through reactivation of TERT transcription {Heidenreich, 2017 #435}. However, in about 15% of cancer cells, the alternative lengthening of telomeres (ALT) pathway can instead support telomere maintenance, a process accomplished by homology-directed repair (HDR) {Apte, 2017 #436;Cesare, 2010 #35}. The ability of cells to adapt mechanisms to maintain their telomeres through recombination was first discovered in budding yeast {Lundblad, 1993 #39}. In fission yeast lacking the telomerase-recruitment factor Ccq1, telomeres gradually shorten with increasing passage number, eventually leading to a growth crisis; the “survivors” that emerge from this crisis can utilize recombination to maintain their telomeres {Tomita, 2008 #1;Miyoshi, 2008 #42}. Fission yeast cells can also survive complete telomere loss by circularization of each of their three chromosomes {Nakamura, 1998 #38}. The generation of these “circular survivors” can occur through multiple pathways, including single-strand annealing, which is independent of Rad51 {Wang, 2008 #437}.

Despite the heterochromatic nature of chromosome ends, telomeres are transcribed into a group of long non-coding RNA species {Azzalin, 2007 #438;Bah, 2012 #365;Bah, 2012 #366;Luke, 2008 #439;Schoeftner, 2008 #440}. The transcription of **TE**lomeric **R**epeat-containing **RNA** (TERRA) is evolutionarily conserved, and has been suggested to function in telomere length regulation, heterochromatin establishment, and telomeric HR {Azzalin, 2015 #441;Bah, 2012 #365;Rippe, 2015 #443}. TERRA transcription is largely RNA Pol II-dependent, is initiated at subtelomeric region(s) and proceeds towards the 3’ end of the telomeric tract {Bah, 2012 #366}. The majority of TERRA molecules are shorter than 500 base pairs (bps) in fission yeast, whereas in mammalian cells TERRA can be as long as several kilobases {Azzalin, 2007 #438;Bah, 2012 #365;Feuerhahn, 2010 #444;Greenwood, 2012 #445;Schoeftner, 2008 #440}. The steady-state level of TERRA is negatively regulated by Taz1 and Rap1 in fission yeast, being very low in wild-type cells {Bah, 2012 #366;Greenwood, 2012 #445}.

Studies in human cells and both budding and fission yeast demonstrate that TERRA molecules localize primarily within the nucleus and partially colocalize with telomeres {Bah, 2012 #366;Cusanelli, 2013 #364;Schoeftner, 2008 #440}. A fraction of TERRA molecules are 3’ polyadenylated {Bah, 2012 #366;Porro, 2014 #446;Schoeftner, 2008 #440}. Poly(A)- and poly(A)+ TERRA have different subnuclear distributions: the majority of poly(A)-TERRA associates with telomeric DNA, whereas poly(A)+ TERRA is mostly released from telomeres and is instead present throughout the nucleoplasm {Moravec, 2016 #423;Porro, 2014 #446}. We recently reported that telomere shortening in fission yeast leads to derepression of TERRA, similar to what has been observed in budding yeast {Cusanelli, 2013 #364;Moravec, 2016 #423}. A substantial fraction of this telomere shortening-induced TERRA is polyadenylated and physically interacts with telomerase, thus promoting its recruitment to the telomere {Moravec, 2016 #423}, potentially in a manner specific to the telomere from which the TERRA was transcribed {Cusanelli, 2013 #364}. Polyadenylation of fission yeast TERRA occurs coincident with 3’ RNA cleavage at sites that lead to loss of nearly the entire G-rich telomere tract {Moravec, 2016 #423}. Therefore, at least in WT fission yeast, only the poly(A)-TERRA carries G-rich telomeric repeat sequences. Taken together, these observations raise the possibility that nascent, telomere-associated TERRA includes the telomeric repeats, but that upon maturation and release from the telomeres, the telomeric-repeat tract is lost. To date the impact of telomere-associated TERRA on telomere maintenance by recombination in the absence of telomerase function in cycling cells has gone largely uninvestigated in fission yeast, although recent evidence suggests that TERRA up-regulation in quiescent fission yeast correlates with increased telomere rearrangements driven by recombination {Maestroni, 2017 #447}.

G-rich TERRA can form RNA-DNA hybrids at the telomere, resulting in a triple-stranded structure called a “telomeric R-loop” (telR-loop); telR-loops are antagonized by the activity of RNase H, which specifically degrades RNA engaged in such hybrids {Balk, 2013 #3;Balk, 2014 #357;Arora, 2014 #448}. Recent evidence indicates that telR-loops may actively engage in HR in telomerase-negative human and yeast cells. For example, depletion of RNase H1 (alone or in combination with RNase H2) results in accumulated RNA-DNA hybrids at telomeres and greater telomere recombination in *cis,* while RNase H over-expression leads to decreased telomere recombination in pre-senescent cells and compromised telomere maintenance in ALT cells {Arora, 2014 #448;Balk, 2013 #3;Balk, 2014 #357}.

In order to test the consequences of TERRA up-regulation, several studies have taken advantage of engineered, inducible promoters at a TERRA transcriptional start site to construct a **t**ranscriptionally **i**nducible **tel**omere (tiTEL) {Maicher, 2012 #449;Pfeiffer, 2012 #450;Arora, 2014 #448;Moravec, 2016 #423}, which can increase telR-loops {Arora, 2014 #448}. However, the strong transcription driven by a heterologous promoter may also disrupt the heterochromatic state of telomeres and influence the maturation, turnover and function of natural TERRA molecules. Despite the growing evidence linking TERRA to regulation of telomeric recombination, it is still not clear how endogenous TERRA (and natively-formed telR-loops) might be altered to promote telomere recombination and formation of telomerase-deficient survivors in an unperturbed system.

Here we study the process by which cells adapt to maintain their telomeres by recombination using a fission yeast model in which Ccq1 is deleted. We chose this genetic system as cells undergo telomere shortening due to defective telomerase recruitment, but are highly proficient at forming linear survivors {Miyoshi, 2008 #42;Tomita, 2008 #1}. Given that Ccq1 also supports repression of subtelomeric heterochromatin involving the SHREC (Snf2/HDAC-containing repressor complex) and CLRC (made up of the H3K9 methylase, Clr4, and an E3 ubiquitin ligase complex composed of Cul4, Rik1, Raf1, and Raf2) complexes {Armstrong, 2018 #408;Sugiyama, 2007 #43;Wang, 2016 #407}, this ability could be linked to Ccq1’s dual roles in telomerase recruitment and subtelomeric silencing. Consistent with this hypothesis, we find that TERRA levels are higher in *ccq1Δ* cells at germination, but dramatically rise further during the emergence of linear, recombination-based survivors. TERRA engages in telR-loops at the telomere(s) and promotes a transition to productive telomere maintenance. Constitutive degradation of RNA-DNA hybrids by over-expression of RNase H drives these linear survivors back into a growth crisis, from which an alternative type of linear survivor arises with distinct cytological features. Surprisingly, the telomeres in these RNase H-resistant survivors no longer recruit Rap1 despite enhanced Taz1 binding. Importantly, deletion of Rap1 insulates linear survivors from the crisis induced by RNA-DNA hybrid degradation, suggesting that this altered telomere composition functionally promotes telomere maintenance when telomere-associated TERRA is degraded.

## MATERIALS AND METHODS

### *S. pombe* strain generation and culture conditions

*S. pombe* strains are described in Table S1. Standard manipulations and cell culturing were carried out as described {Moreno, 1991 #319}. C-terminal GFP or mCherry tagging was performed with the pFa6a-GFP-kanMX6 or pFa6a-mCherry-kanMX6 cassette {Bähler, 1998 #199;Snaith, 2005 #472}. pFa6a-natMX6-nmt41-HA was used to tag Rnh201 at the N-terminus as established {Bähler, 1998 #199}. In the strain with “transcriptionally inducible” telomere (tiTEL), the thiamine repressible *nmt1+* gene promoter (*Pnmt1*) was inserted 91 base pairs upstream of the endogenous TERRA transcription start site at Chr I-R {Moravec, 2016 #423}. A C-terminal GFP cassette was further integrated after the Taz1 ORF in this tiTEL strain (Strain MKSP1722). Fresh pre-senescent cells for growth assays of *ccq1*Δ, *trt1*Δ, *rad55*Δ, and combinations or variants of these alleles were obtained by dissection of sporulated heterozygous diploid strains. In the case of h+/h+ diploids, strains were first transformed with plasmid pON177 {Styrkarsdottir, 1993 #469} to provide the mat-M genes. All strains generated by cassette integration were confirmed by PCR. After genetic crosses, progeny were tested by the segregation of markers, PCR, or presence of the relevant fluorescent protein fusion, as appropriate.

*S. pombe* were grown at 30°C, in yeast extract medium supplemented with amino acids (YE5S) or Edinburgh minimal medium with amino acids (EMM5S). To suppress the *nmt1+* or *nmt41+* gene promoter {Maundrell, 1990 #201}, 5 μg/ml thiamine (Sigma-Aldrich) was added to EMM5S medium.

### Serial culturing

Single colonies formed from fresh dissections were inoculated in liquid medium. 24 hours later, cell densities were measured using the Moxi-Z mini cell counter (ORFLO). Approximately 28 generations were estimated from a single, germinated cell growing on solid medium for 20 generations to generate a colony, followed by a single overnight culture. Cell cultures were measured for their cell densities and diluted to 5×10^5^ cells/ml with fresh medium every 24 hours. At each indicated time point, cells were collected for further analysis or subjected to imaging as described. The increase in generations each day were calculated as log_2_([cell density]/(5×10^5^)). “Survivors” indicate that the culture has achieved a consistent growth rate, typically around 200 generations (> 20 days). Growth curves were plotted using GraphPad Prism 7.01.

### Live cell imaging and analysis

Cells for imaging were exponentially grown in YE5S or EMM5S media supplemented with adenine hemi sulfate (0.25 mg/mL). Cells in liquid culture were concentrated and applied to glass slides bearing agarose pads (1.4% agarose in EMM). Samples were sealed under a glass coverslip with VALAP (equal parts by weight: Vaseline, lanolin, and paraffin). Images were acquired on an Applied Precision (now GE Healthcare) Deltavision high performance wide field microscope with solid-state illumination and a Photometrics Evolve 512 EMCCD camera. For imaging of Taz1-GFP in Fig. 4A and Fig. S3A-C, each field was captured as a 12 section Z-stack of sample thickness 4.80 μm (optical sectioning every 0.40 μm; this is sufficient to capture the entire nuclear volume to ensure all telomeric foci are visualized); EMCCD Gain = 100; GFP at 10% power, 100 ms exposure. For telomere volume reconstruction (Fig. 4D), each field was captured as a 25 section Z-stack of sample thickness 5.0 μm (optical sectioning spacing every 0.2 μm); EMCCD Gain = 100; GFP at 10% power, 50 ms exposure. For cells expressing Rap1-GFP (Fig. 5A,B; Fig. S3C,D) each field was captured as a 24 section Z-stack of sample thickness 4.80 μm (optical sectioning every 0. 2 μm; EMCCD Gain = 200; GFP at 10% power, 25 ms exposure. For visualization of Taz1-GFP and Rad52-mCherry (Fig. 1B,C) each field was captured as a 12 section Z-stack of sample thickness 4.8 μm (optical sectioning spacing every 0.4 μm); EMCCD Gain = 100; GFP at 10% power, 50 ms exposure; mCherry at 10% power, 500 ms exposure. Only live, non-mitotic cells that had completed cytokinesis were considered for Taz1-GFP and cell length analysis. In most cases we display maximum intensity projections, as described in the Legends.

**Figure 1.**
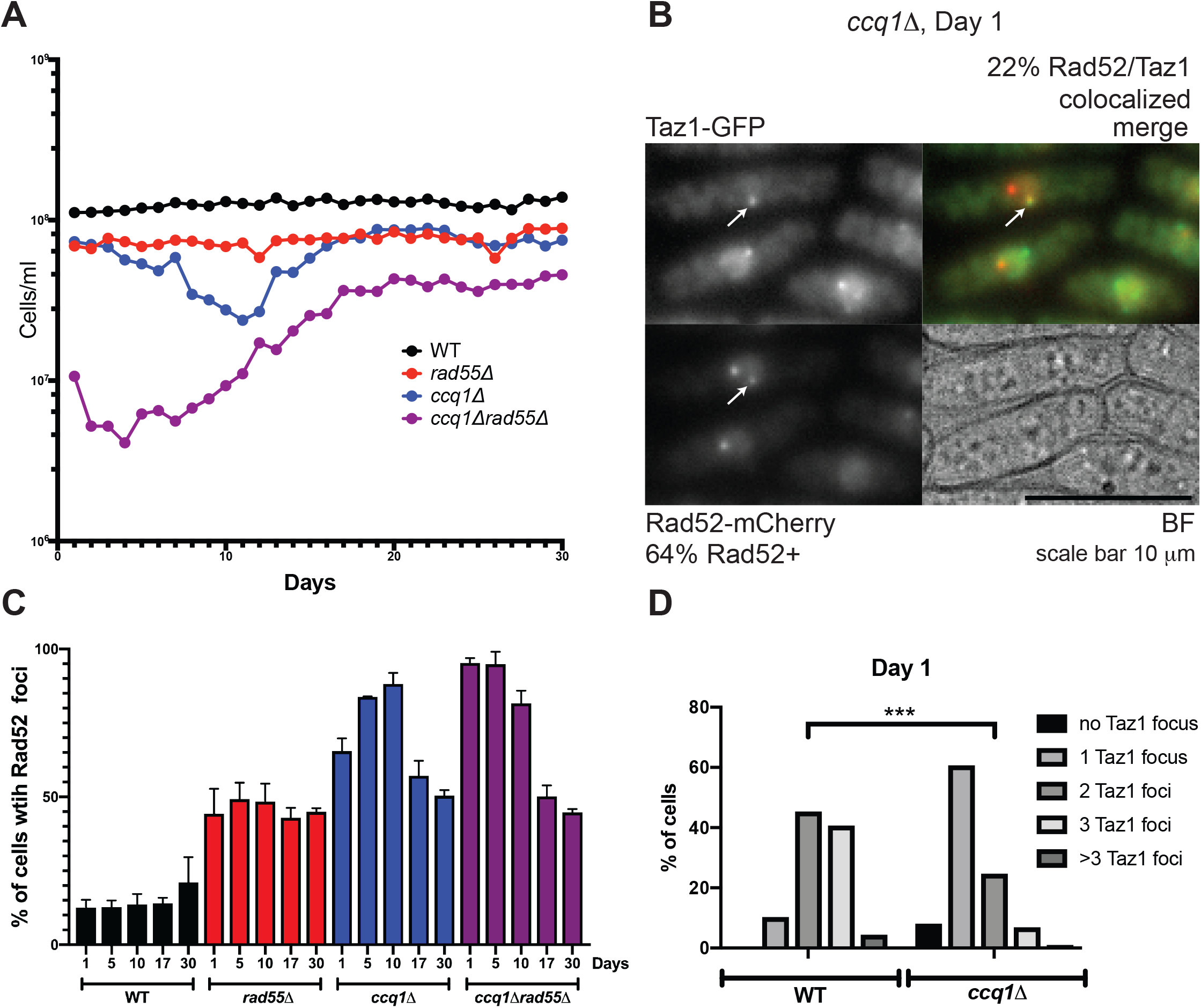
Recombination factors are loaded at telomeres shortly after germination. (A) Serial culturing of WT, *ccq1Δ, rad55Δ* and *ccq1Δrad55Δ* cells. Plot of cell density versus number of days in culture since germination, diluted once every 24 hours. (B) Telomeres are recognized as DNA damage shortly after germination in *ccq1Δ* cultures. Representative micrographs of *ccq1Δ* cells expressing Taz1-GFP and Rad52-mCherry immediately after germination. The percent of cells with the indicated phenotype are noted. BF = brightfield; Scale Bar = 10 μm (C) The percent of cells with Rad52-GFP foci mirrors growth rate, with *ccq1Δrad55Δ* cells displaying high levels of Rad52 foci at germination. WT, *ccq1Δ, rad55Δ* and *ccq1Δrad55Δ* cells expressing Rad52-GFP were imaged on the indicated days of the growth assay in (A). n>100 cells for each strain and timepoint from at least 3 biological replicates. Images displayed are maximum intensity projections of Z-stacks obtained as described in the Methods. (D) The number of Taz1-GFP foci is less in *ccq1Δ* cells than WT cells on Day 1. p < 0.001 by K-S test.

Images were analyzed in ImageJ (Fiji). To calculate the number of Taz1-GFP foci per cell or examine the co-localization of Taz1-GFP and Rad52-mCherry, maximum Z-projections were generated. For quantitative measurement of Taz1-GFP recruitment to telomeres, the intensity of each Taz1-GFP focus was measured from mean intensity projections as the integrated pixel density of the visibly occupied area. The background GFP signal was removed from measurement of each focus by subtracting the integrated density of an empty area of the same size as the focus within the nucleoplasm of the same cell. The intensities of all foci were then summed to give a single measurement per cell.

### Real-time PCR to measure TERRA levels

RT-PCR protocols were adapted from our previous work {Bah, 2012 #366}. Total RNA was extracted using MasterPure yeast RNA purification kit (Epicentre). To completely eliminate DNA contamination, RNA was treated three times with RNase-free DNase I (provided with the kit). cDNA was generated using SuperScript III (Invitrogen). Each 20 μl reverse transcription reaction contained 5 μg total RNA, 1 μl 10 μM oligo for total TERRA (Sp C-rich telo: 5’-TGTAACCGTGTAACCACGTAACCTTGTAACCC-3’) and 1 μl 1 μM reverse control oligo for actin (Sp ACT1(2): 5’-CATCTGTTGGAAAGTAGAAAGAGA AGCAAG-3’), or 2 μl 40 μM oligo(dT) (in the case of poly(A)+ TERRA), 1 μl 10 mM dNTP, 4 μl 5xFirst-Strand Buffer, 2 μl 0.1 M DTT and 1 μl SuperScript III RT enzyme. The reaction was incubated at 50°C for 1 hour and the remaining RNA was degraded by RNase H (NEB) and RNase A (Sigma-Aldrich) by incubating at 37°C for 20 minutes. Real-time PCR was performed on C1000 Touch Thermal Cycler (Bio-Rad), using FastStart Universal SYBR Green Master (Rox) (Roche). For TErRa pCr amplification, each 20 μl reaction contained 2 μl of cDNa, 1 μl of each 10 μM primer (SpRACE290-320_4: 5’-GGGCCCAATAGTGGGGGCATTGTATTTGTG-3’; Sp-subtelo_425: 5’-GAAGTTCACTCAGTCATAATTAATTGGGTAACGGAG-3’). For actin PCR amplification, each 20 μl reaction contained 2 μl of 1:10 diluted cDNA, 1 μl of each 10 μM primer (SpACT1FOR2: 5’-CCGGTATTCATGAGGCTACT-3’; SpACT1REV2: 5’-GGAGGAGCAACAATCTTGAC-3’).

qPCR steps: 1 cycle of 95°C-5 min; 45 cycles of 95°C-10 sec and 60°C-30 sec followed by a melting curve. Relative TERRA levels were calculated for each sample after normalization against actin using the ΔΔCt method. Ct values for each reaction were determined using the software Bio-Rad CFX Manager 2.1. Duplicate Ct values were averaged in each run. Each RNA sample was reverse-transcribed into cDNA twice, and each cDNA sample was analyzed by RT-PCR at least three times. Ct values of the same sample were averaged and plotted in GraphPad Prism 6. No RT and no template reactions were always included to control for DNA contamination.

### Southern blots

To measure telomere length, experiments were performed as described {Bennett, 2016 #473}. Briefly, genomic DNA (25 μg) was digested with EcoRI-HF (NEB) at 37°C overnight, electrophoresed on a 1% agarose gel, denatured and transferred to Zeta-Probe GT membrane (Bio-Rad), then hybridized with a telomere-DIG probe in ExpressHyb Solution (Clontech). After hybridization, the membrane was washed and incubated with anti-DIG-AP conjugate (the same kit, 1:1000 dilution). The blot was then treated with SuperSignal West Femto Maximum Sensitivity Kit (ThermoFisher/Pierce), and imaged on VerSaDoc imaging system (Bio-Rad) using the software Quantity One 4.6.9.

### Pulsed-field gel electrophoresis

Experiments were performed as in {Ferreira, 2001 #421;Nakamura, 1998 #38} with minor modifications described in {Lorenzi, 2015 #422}. Briefly, cells were washed in TSE (10 mM Tris–HCl (pH 7.5), 0.9 M D-Sorbitol, 45 mM EDTA) and resuspended at 5 × 10^7^ cells with 10 mg/ml zymolyase-20T (Seikagaku Biobusiness) and 50 mg/ml lysing enzyme (Sigma). One volume of 2% low melt agarose (Bio-Rad) in TSE was added and the suspension dispensed into plug molds. Plugs were incubated in TSE containing 5 mg/ml zymolyase-20T and 25 mg/ml lysing enzyme at 37°C for 1 h. Plugs were incubated for 90 min at 50°C in 0.25 M EDTA, 50 mM Tris–HCl (pH 7.5), 1% SDS and then for 3 days at 50°C in 0.5 M EDTA, 10 mM Tris–HCl (pH 9.5), 1% lauroyl sarcosine, 1 mg/ml proteinase K. Plugs were washed in 10 mM Tris–HCl (pH 7.5), 10 mM EDTA and incubated with 0.04 mg/ml PMSF for 1 h at 50°C. Plugs were washed again in 10 mM Tris–HCl (pH 7.5), 10 mM EDTA and then digested with 100 U of *NotI* (New England Biolabs) for 24 h at 37°C. Plugs were equilibrated in 0.5× TAE, loaded on a 1% agarose gel, and ran in a CHEF-DR III system (Bio-Rad) for 16 h at 14°C using the following program: 60-120 s switch time, 120° angle, 6 V/cm electric field. After running, gels were processed as for Southern blot and hybridized with a mix of L, I, M, and C probes recognizing unique sequences from the most terminal *NotI* fragments of the left arm of chromosome I, right arm of chromosome I, left arm of chromosome II, and right arm of chromosome II, respectively. For Southern blotting, DNA was denatured in gel and transferred to a positively charged nylon membrane (GE Osmonics) and processed as described above.

### Chromatin Immunoprecipitation (ChIP)

ChIP of Rap1-GFP expressing cells was carried out as described previously {Lorenzi, 2015 #422} with the following modifications. 75 OD of yeast cells at an of OD_600_ 0.6–1.0 were cross-linked in 1% formaldehyde for 30 min prior to quenching with glycine. Cross-linked material was washed in ice-cold PBS and resuspended in 1ml lysis buffer (50 mM HEPES-KOH pH 7.5, 140 mM NaCl, 1 mM EDTA, 1% Triton X-100, 0.1% sodium deoxycholate, protease inhibitor cocktail (Roche)). Cells were transferred to 1.5 ml screw-cap tubes and subjected to mechanical lysis with silica beads (0.5 mm diameter, BioSpec) using the Mini-BeadBeater-16 instrument (BioSpec), shaking 1 min 30 sec for 6 – 8 times with 3 min intervals. Lysates were centrifuged for 30 min at 16,000xg, and pellets were resuspended in 500 μl lysis buffer and subjected to sonication using the S-4000 sonicator (Misonix) for 15 min (Amplitude 4, 5 sec pulse with 15 sec cool-down intervals). After centrifugation at 10,000xg for 15 min, the supernatant containing DNA fragments was collected. Lysis buffer with protease inhibitor cocktail (Roche) was added to adjust the volume to about 1.4 ml with 120 μl taken as the input sample. Immunoprecipitations were performed on a rotating wheel at 4°C overnight with GFP-Trap beads (Chromotek). The beads were then washed 4 times with lysis buffer, lysis buffer plus 500 mM NaCl, wash buffer (10 mM Tris–HCl pH 7.5, 0.25 M LiCl, 0.5% NP40, 0.5% sodium deoxycholate), and lysis buffer respectively. Immunoprecipitated chromatin was eluted by adding 120 μl elution buffer (1% SDS, 100 mM NaHCO_3_, 40 ug/ml RNase A) and incubated at 37°C for 1 h. The input sample was treated in parallel by adding 6 μl of 20% SDS, 11 μl of 1 M NaHCO_3_ and RNase A. Cross-links were reversed at 65°C overnight (16 hrs) and DNA was purified using a PCR purification kit (Qiagen). DNA content was quantified by RT-PCR using iTaq Universal SYBR Green Supermix (Bio-Rad) on a C1000 Thermal Cycler instrument (Bio-Rad) with primers: subtel-for (5’-TATTTCTTTATTCAACTTACCGCACTTC-3’) and subtel-rev (5’-CAGTAGTGCAGTGTATTATGATAATTAAAATGG-3’).

### Western blots

Experiments were carried out as described {Lorenzi, 2015 #422}. Cells were collected at exponential phase and samples were prepared by TCA precipitation. Antibodies were as follows: a rabbit polyclonal anti-Rap1 and a rabbit polyclonal anti-Taz1 antibody (kind gifts from J. Kanoh and J. P. Cooper, respectively); and a mouse monoclonal anti-beta actin antibody (mAbcam 8224) used as loading control.

### DNA immunoprecipitation

Log phase cells were harvested, washed with water, and flash frozen in liquid nitrogen. The frozen pellet was resuspended in 1 ml of RA1 buffer (Macherey-Nagel) supplemented 1% β-mercaptoethanol. The cell suspension was mixed with 300 μl of phenol:chloroform:isoamyl alcohol (25:24:1 saturated with 10 mM Tris–Cl pH 8.0, 1 mM EDTA) and 100 μl of acid washed glass beads. Cells were lysed by mechanical shaking using a cell disruptor (Disruptor Gene) for 3 minutes using the 30 second on/off mode at power 4. Cell extracts were centrifuged (13,000g, 20 min, 4 °C). Pellets were washed with 70% ethanol, air-dried, and re-suspended in 200 μl of Tris–EDTA and sonicated with a Bioruptor (Diagenode) to obtain 100–500 bp long fragments. Ten micrograms of sheared nucleic acids were diluted in 1 ml of IP buffer (0.1% SDS, 1% Triton X-100, 10 mM HEPES pH 7.7, 0.1% sodium deoxycholate, 275 mM NaCl) and incubated overnight on a rotating wheel at 4 °C in presence of 1 μg of S9.6 antibody (Kerafast) and 20 μl of protein G sepharose beads (GE Healthcare) blocked with *Escherichia coli* DNA and bovine serum albumin. Beads were washed four times with IP buffer and bound nucleic acids were isolated using the Wizard SV gel and PCR clean-up kit (Promega). Collected DNA fragments were quantified using RT-PCR as described above with the following primers: oF1 (5’-GAAGTTCACTCAGTCATAATTAATTGGGTAACGGAG-3’) and oR1 (5’-GGGCCCAATAGTGGGGGCATTGTATTTGTG-3’).

### Data Availability Statement

Table S1 describes all fission yeast strains used in this work, and are available upon request. The authors affirm that all data necessary for confirming the conclusions of the article are present within the article, figures, and tables.

## RESULTS

### Telomere recombination likely precedes the growth crisis in *ccq1Δ* cells

Immediately following germination, *ccq1Δ* cells propagated in liquid culture grow slightly slower than WT cells (Figure 1A and Figure S1A) but already have substantially shorter telomeres as measured by Southern blot (Figure S1B). After ten days of serial culturing, *ccq1Δ* cells are in the depth of the growth crisis, which is followed by the emergence of survivors that have growth rates similar to the starting population, consistent with earlier studies {Flory, 2004 #409;Harland, 2014 #396;Miyoshi, 2008 #42;Tomita, 2008 #1} (Figure 1A and Figure S1A). These observations suggest that pre-senescent *ccq1Δ* cells have already undergone substantial telomere loss prior to entry into the growth crisis (Figure 1A, Figure S1A,B), consistent with previous studies in budding yeast {Khadaroo, 2009 #452;Abdallah, 2009 #453} and with the activation of the ATR-dependent Chk1 checkpoint {Tomita, 2008 #1} and formation of Crb2 foci {Carneiro, 2010 #2} in early generation *ccq1Δ* cells. In light of these results, we considered whether telomere recombination was occurring shortly after germination, but was insufficient to maintain telomere length. To assess this, we serially cultured cells lacking both Ccq1 and Rad55/Rhp55, which ultimately survive primarily through chromosome circularization {Miyoshi, 2008 #42}. Consistent with our hypothesis and the observed role for the recombination factor Rad52 in antagonizing senescence in budding yeast models of telomere dysfunction {Abdallah, 2009 #453;Lundblad, 1993 #39}, *ccq1Δ/rad55Δ* cells grow much more poorly than *ccq1Δ* cells immediately following germination (Figure 1A and Figure S1A). This suggests that the activation of recombination at critically short or deprotected telomeres occurs early after germination of *ccq1Δ* cells but is insufficient to maintain telomere length.

Shortly after germination we also observe Rad52(Rad22)-mCherry foci in ~65% of *ccq1Δ* cells (n=233) compared to only 2% of WT cells (n=239) (Figure 1B,C). While only a subset of these foci colocalize with Taz1-GFP (22% of *ccq1Δ* cells; Figure 1B), it is likely that the remaining Rad52-mCherry foci correspond to critically short telomeres that cannot recruit sufficient Taz1 to be visualized. Indeed, the percentage of cells with 2-3 Taz1-GFP foci decreases from about 85% for WT cells to only 30% of *ccq1Δ* cells (Figure 1D). Nearly all cells lacking both Ccq1 and Rad55 have Rad52-GFP foci at this initial time point (Figure 1C), further suggesting that Rad52 foci are tied to telomere dysfunction. Importantly, cells lacking Rad55 alone grow more slowly than WT (Figure 1A) and have consistent, persistent Rad52-GFP foci (Figure 1C) over the course of the serial culturing assay rather than the dynamic increase and subsequent decrease in Rad52-mCherry foci seen during telomere crisis and escape. Taken together, these results suggest that recombination is active at *ccq1Δ* telomeres shortly after germination but that additional factors must underlie the ability of cells to productively use recombination to maintain their telomeres, as seen in survivors.

### TERRA levels increase in recombination-based survivors, but driving TERRA production cannot prevent telomere crisis in pre-senescent *ccq1Δ* cells

Studies in both budding yeast and human cells suggest that TERRA up-regulation underlies productive recombination to maintain telomeres in the absence of telomerase, which correlates with elevated levels of telR-loops {Arora, 2012 #363;Balk, 2013 #3;Graf, 2017 #413;Yu, 2014 #454}. Consistent with the role for Ccq1 in maintaining subtelomeric heterochromatin {Armstrong, 2018 #408;Sugiyama, 2007 #43;Wang, 2016 #407}, we find that TERRA levels, which are already ~10-fold higher in *ccq1Δ* cells shortly after germination than WT cells, are dramatically increased in *ccq1Δ* survivors irrespective of whether Taz1-GFP, a convenient cytological marker for telomeres, was expressed during serial culturing (Figure 2A). Indeed, we observe a progressive increase in TERRA during the emergence of *ccq1Δ* survivors as detected by quantitative RT-PCR (Figure 2B). Moreover, we detect an increase in telR-loops by DNA:RNA-IP/RT-PCR in *ccq1Δ* survivors (Figure 2C). We were also curious whether TERRA levels increase in survivors derived from other genetic backgrounds. Consistent with the ability of TERRA to support ongoing telomeric recombination in budding yeast {Balk, 2013 #3}, we observe mild TERRA induction in recombination-competent *trt1Δ* survivors, which when serially cultured in liquid media likely contain both linear and circular types of survivors {Nakamura, 1998 #38} (Figure S1C). By contrast, although *ccq1Δ/rad55Δ* cells show very high levels of TERRA shortly after germination (at 38 generations), *ccq1Δ/rad55Δ* survivors have nearly undetectable TERRA levels, far below what is seen in WT cells, likely due to complete loss of telomeric repeats (Figure 2D). Taken together, these results show that TERRA production is quickly stimulated in cells lacking Ccq1 and is either further increased during formation of survivors (in *ccq1Δ* recombination-competent strains) or is lost (in recombination-deficient genetic backgrounds that undergo chromosome circularization such as *ccq1Δ/rad55Δ).*

**Figure 2.**
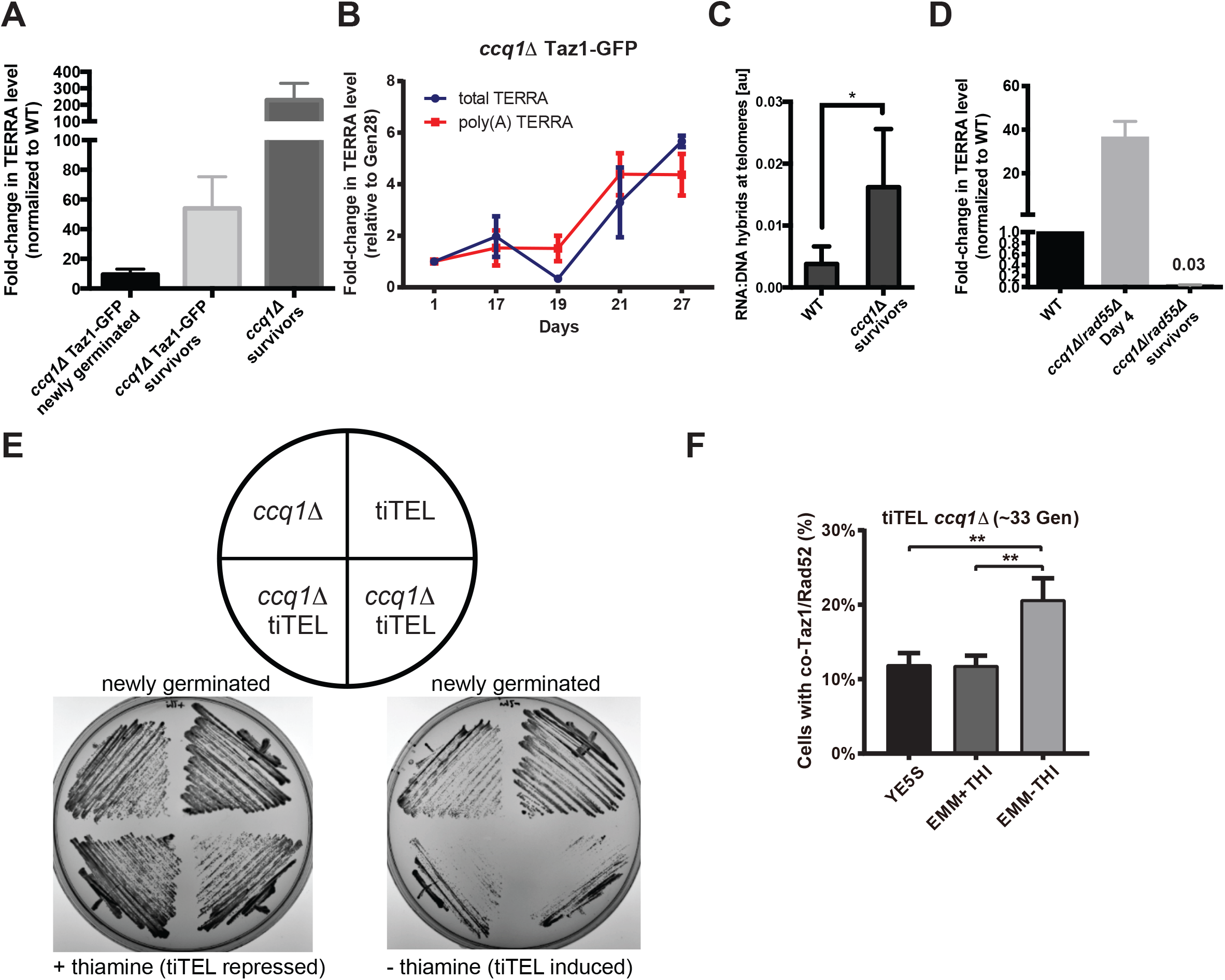
TERRA is up-regulated during *ccq1Δ* survivor formation but a tiTEL cannot prevent a growth crisis. (A) TERRA levels are higher in *ccq1Δ* cells shortly after germination and are dramatically increased in survivors. Quantitative RT-PCR analysis for G-rich TERRA normalized to WT cells and plotted as the mean with SD from three replicates. (B) TERRA levels increase concomitant with the emergence of *ccq1Δ* survivors between Days 19-27. Quantitative RT-PCR analysis for both total and poly(A+) TERRA normalized to Day1 *ccq1Δ* cells and plotted as the mean with SD from three replicates. (C) Telomeric RNA/DNA hybrids increase in *ccq1D* survivor cells as assessed by DNA-IP using the S9.6 antibody (see Methods). Mean of three replicates plotted with the standard deviation. * = p < 0.05. (D) Newly germinated *ccq1Δrad55Δ* cells contain high levels of total TERRA shortly after germination but TERRA is depleted in *ccq1Δrad55Δ* survivors as determined by RT-PCR as in (A). (E,F) tiTEL induction leads to a more rapid and pronounced growth crisis in newly germinated *ccq1Δ* cells. (E) Culturing of two *ccq1Δ* tiTEL clones on either EMM + thiamine (tiTEL repressed) or – thiamine (tiTEL induced) solid media. tiTEL induction leads to poorer growth than seen for *ccq1Δ* cells alone. (F) Increased loading of recombination factors on telomeres in newly germinated *ccq1Δ* tiTEL cells upon induction. Fluorescence micrographs of newly germinated (33 generations) ccq1Δ tiTEL cells expressing Taz1-GFP and Rad52-mCherry were quantified for co-localization. ** p<0.01 by Student’s t-test.

Our data suggest that TERRA up-regulation during the emergence of survivors from the growth crisis in *ccq1Δ* cells supports recombination-based telomere maintenance. However, we were curious whether driving TERRA production from the time of germination would be sufficient to avoid the growth crisis in *ccq1Δ* cells altogether. To examine this, we took advantage of the tiTEL system in which the thiamine-regulated *NMT1* promoter drives TERRA over-expression from a single telomere {Moravec, 2016 #423}. Surprisingly, *ccq1Δ* tiTEL cells serially restruck on plates under conditions of TERRA induction grow much more slowly than *ccq1Δ* cells or tiTEL cells grown under the same conditions (Figure 2E). Interestingly, we find that Rad52-mCherry is enriched on telomeres one day after tiTEL induction (but not in cells in which the tiTEL was repressed), consistent with signals that slow cell growth (Figure 2F). Thus, engineered TERRA over-expression cannot overcome the passage of cells through a period of catastrophic growth crisis.

### Degradation of RNA-DNA hybrids drives *ccq1Δ* survivors to enter a second growth crisis from which new survivors emerge

To test whether TERRA up-regulation (and specifically TERRA engaged in telR-loops) contributes to telomere maintenance by recombination in *ccq1Δ* cells, we sought to develop an approach to induce degradation of telR-loops. To this end, we integrated the inducible *nmt41* promoter in front of the Rnh201 gene, which encodes an RNase H2 homologue capable of digesting RNA specifically engaged in RNA-DNA hybrids and TERRA in budding yeast when overexpressed {Balk, 2013 #3;El Hage, 2010 #327;Huertas, 2003 #328}.

We found that Rnh201 over-expression (Figure S2A-B) is sufficient to suppress telR-loops in S. *pombe* (Figure S2C), although we note that even in the repressed state (with thiamine) Rnh201 is over-expressed relative to WT cells (Fig. S2B). Strikingly, overexpression of Rnh201 caused *ccq1Δ* survivors to rapidly reenter a growth crisis (Figure 3A), characterized by cell cycle arrest as indicated by the accumulation of very long cells (Figure 3B). We also observed hallmarks of telomere crisis 24 hours after induction of Rnh201 over-expression in *ccq1Δ* survivors, including a greater than 3-fold increase in the percent of cells with Rad52 foci (to >60% of all cells), with 80% of these Rad52 foci occurring at the telomere as assessed by colocalization with Taz1-GFP (Figure 3C and Figure S2D). In addition, Rnh201 over-expression in *ccq1Δ* survivors leads to an increase in cells with anaphase bridges decorated with Rad52, suggestive of fused or entangled telomeres due to the ongoing crisis (Figure 3C and Figure S2D). Ultimately “new” survivors emerged about 50 generations after Rnh201 induction (Figure 3A) despite persistent expression of Rnh201 (Figure S2E). These “new” survivor cells, which we term RNase H2-resistant (RHR) *ccq1Δ* survivors, maintain their telomere length as assessed by Southern blot (Figure 3D). Moreover, pulsed-field gel electrophoresis demonstrates that, like the initial *ccq1Δ* survivors, RHR *ccq1Δ* survivors maintain linear chromosomes (Figure 3E) without any apparent telomere-telomere fusions (M+C and I+L bands) that appear in circular *trt1Δ* and *ccq1Δrad55Δ* survivors.

**Figure 3.**
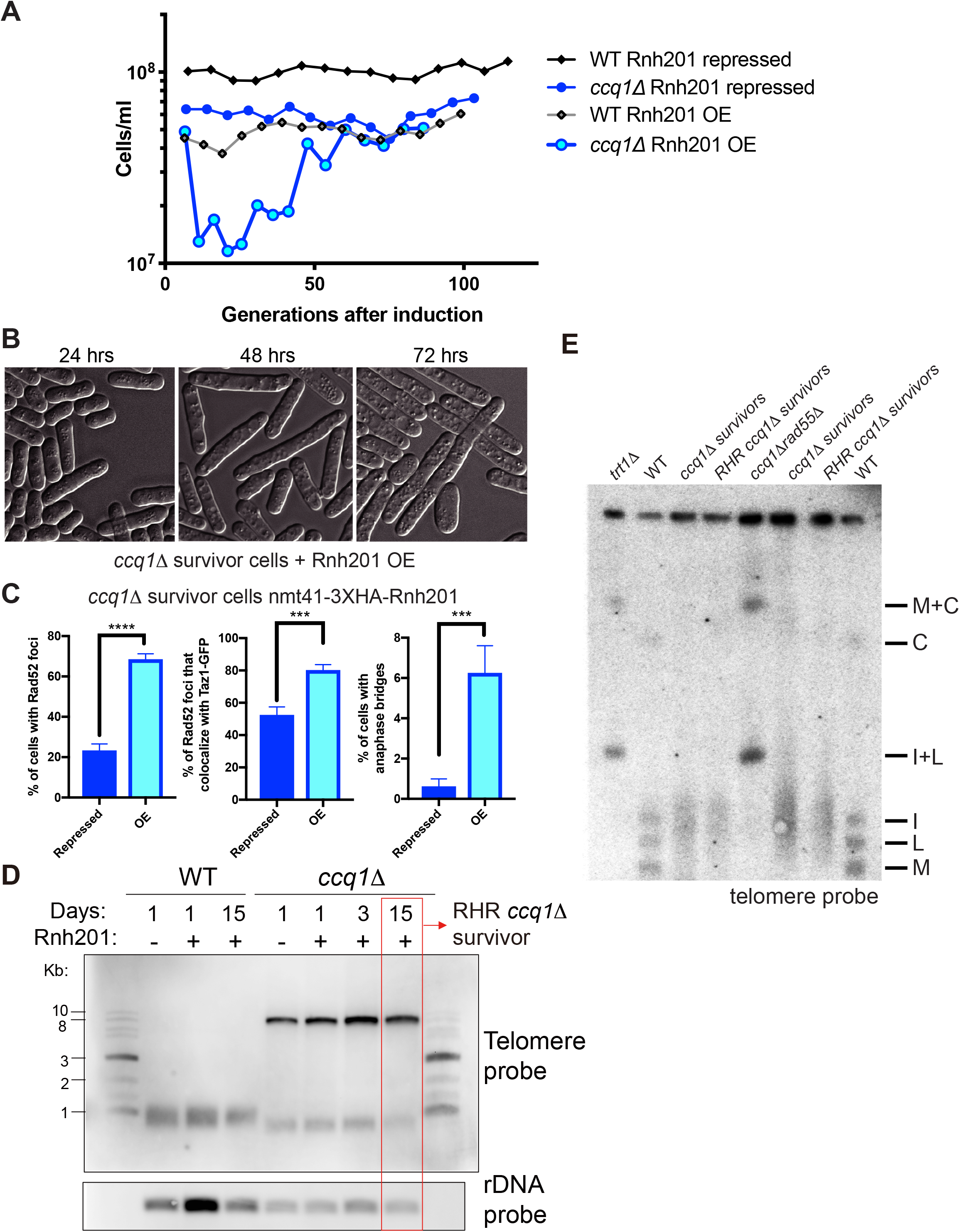
Degradation of RNA-DNA hybrids induces a second growth crisis in *ccq1Δ* survivors, ultimately leading to a new type of RNase H-resistant *ccq1Δ* survivor. (A) Rnh201 induction returns *ccq1Δ* survivors into a growth crisis. WT or *ccq1Δ* Taz1-GFP survivor cells (MKSP1213 and MKSP1214) were modified to insert the nmt41 promoter upstream of the *Rnh201* coding sequence. Strains were then cultured in a repressed state (with thiamine) or induced state (without thiamine), as indicated and diluted every 24 hours. (B) DIC imaging shows that Rnh201 over-expression leads to a new growth crisis in *ccq1Δ* survivors. The first three days of liquid culturing corresponding to Figure 3A are shown. Note the increase in cell length, indicating checkpoint arrest. (C,D) The telomeres of RHR *ccq1Δ* survivors selected after constitutive Rnh201 over-expression are similar in length and structure to the initial *ccq1Δ* survivors. Southern blot of EcoRI digested genomic DNA as in Fig. S1B. The rDNA probe serves as a loading control. (D) The RHR *ccq1Δ* survivors, like *ccq1Δ* survivors, retain linear chromosomes as assessed by pulsed-field gel electrophoresis followed by Southern blotting. Genomic DNA was digested with *NotI* and hybridized to C, I, L, and M probes, which detect the terminal fragments of chromosomes I and II. Bands corresponding to chromosome end fusions are indicated by M+C and I+L bands, observed in the *trt1Δ* and *ccq1Δrad55Δ* circular survivors.

### RHR *ccq1Δ* survivors display altered recruitment of telomere binding proteins and compartmentalization of telomeres

To gain further insights into the mechanisms that RHR *ccq1Δ* survivors employ to maintain linear chromosomes, we visualized telomeres by expression of Taz1-GFP. Surprisingly, Taz1-GFP intensity in the initial *ccq1Δ* survivors increased compared to WT cells (Figure 4A,B). In addition, some *ccq1Δ* survivors have a single, very bright Taz1-GFP focus (Fig. 4A, center panel, arrowhead). The generation of RHR *ccq1Δ* survivors led to an even higher Taz1-GFP intensity and more declustered, expanded telomere foci (Figure 4A,B). To further quantitate this defect, we took advantage of an image analysis pipeline we initially developed to render and measure the size of heterochromatin foci in fission yeast {Schreiner, 2015 #474}. In WT cells, the true size of telomere clusters is nearly always smaller than the diffraction limit of our microscope, and thus they appear equivalent to the point spread function (PSF; Figure 4C,D). The mean size of the Taz1-associated telomere clusters in the initial *ccq1Δ* survivors is greater than the PSF (Figure 4C,D). However, the telomere foci in the RHR *ccq1Δ* survivors are highly expanded (Figure 4A-C) and irregularly shaped (Figure 4D). These effects were not influenced by Taz1-GFP expression during the serial culturing, as integration of Taz1-GFP into *ccq1Δ* and RHR *ccq1Δ* survivors led to cytologically indistinguishable telomere appearance (Figure S3A,B).

**Figure 4.**
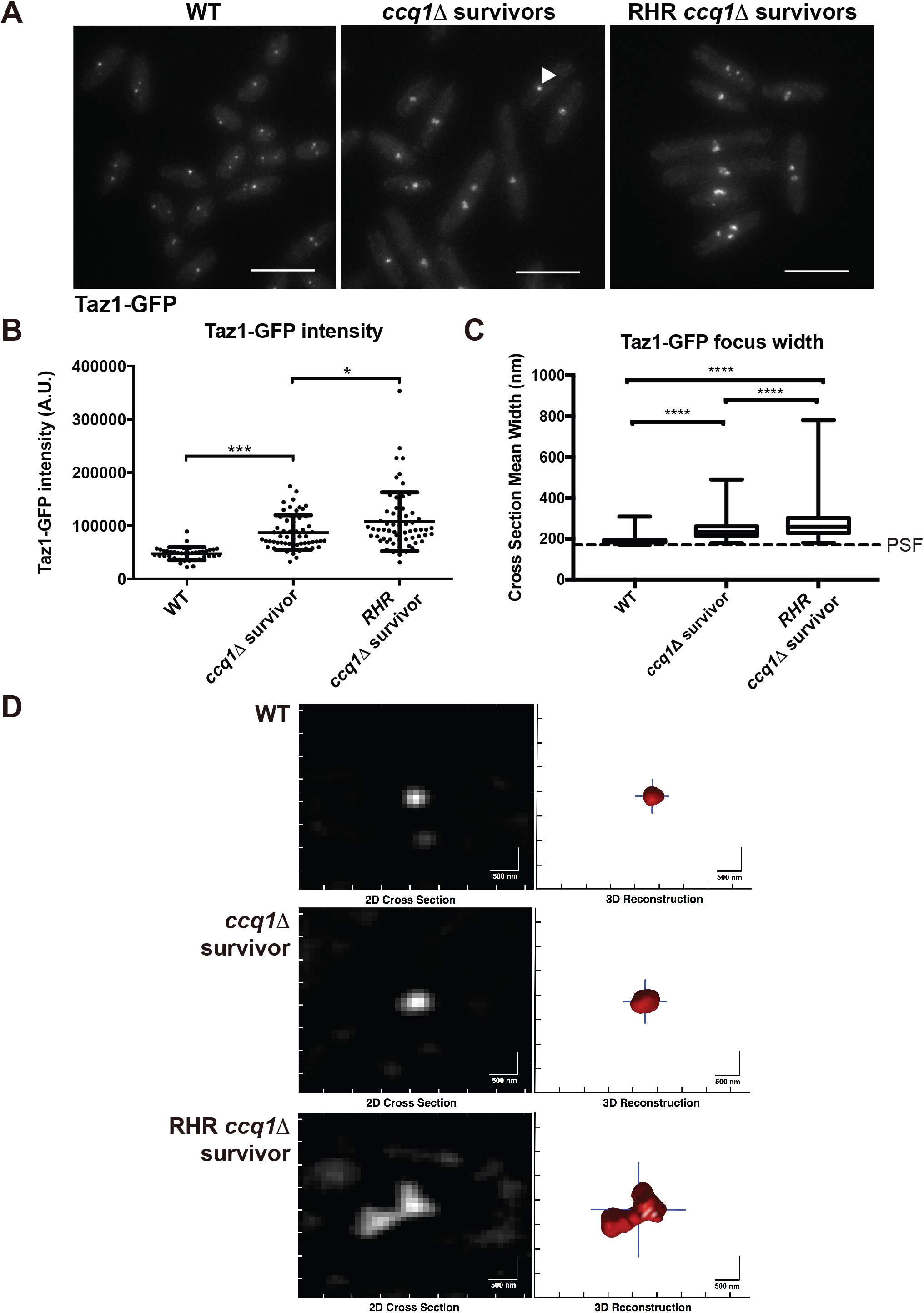
*ccq1Δ* survivors that form after constitutive Rnh201 over-expression display increased Taz1-GFP recruitment and have telomeres that appear larger and declustered. (A) Fluorescence imaging of Taz1-GFP reveals changes in telomere appearance in *ccq1Δ* Taz1-GFP survivors and RHR *ccq1Δ* Taz1-GFP survivor cells. Maximum intensity projections of fluorescence micrographs of WT Taz1-GFP, *ccq1Δ* Taz1-GFP survivors or RHR *ccq1Δ* Taz1-GFP survivor cells obtained as described in the Methods. (B) The intensity of focal Taz1-GFP increases in *ccq1Δ* survivors, and is further enhanced in RHR ccq1Δ survivors. Measurement is the sum intensity of all Taz1-GFP foci. n>50 cells for each genotype. * p<0.05, *** p<0.001 by Student’s t-test corrected for multiple comparisons. (C) Taz1-GFP foci were reconstructed using an algorithm previously developed by our group (see Methods) on Z-stacks obtained as in (A). The mean cross-section width was determined for each focus; for foci with a crosssection smaller then the point spread function (PSF), the focus will appear with the dimensions of the PSF. Reconstructed WT Taz1-GFP foci (n=352) are nearly all the size of the PSF (170 nm, dashed line) or smaller. While *ccq1Δ* survivors display slightly larger Taz1-GFP foci (n=252), the largest Taz1-GFP foci are seen in the RHR *ccq1Δ* survivors (n=132). The box represents the 25^th^ and 75^th^ quartiles, the mean is the black line and the whiskers include all data. **** p < 0.0001 by Student’s t-test corrected for multiple comparisons. (D) Representative examples of reconstructed Taz1-GFP foci in the three genetic backgrounds. The raw image is on the left and the telomeric focus is reconstructed on the right. Scale bar = 500 nm.

In budding yeast, telomere clustering is promoted by the shelterin component, Rap1, acting in concert with Sir3 {Gotta, 1996 #455;Hoze, 2013 #456;Ruault, 2011 #457}. Loss of Rap1 alone is insufficient to cause telomere declustering in fission yeast (Figure S3C) and Rap1 foci appear unaltered in *ccq1Δ* survivors (Figure S3D,E). However, we observe a profound loss of Rap1-GFP from RHR *ccq1Δ* survivor telomeres (Figure 5A) despite normal (or higher) levels of Taz1-GFP recruitment (Figure 4). Indeed, in 45% of cells we could not detect any Rap1-GFP at the telomere, while 2 or more Rap1-GFP-associated telomere clusters were seen only very rarely (Figure 5B). Chromatin immunoprecipitation followed by qPCR further supported this observation, as we find a significant decrease in the association of Rap1-GFP with the subtelomere upon longterm Rnh201 over-expression in *ccq1Δ* survivors (Figure 5C), although we acknowledge that loss of Rap1 binding could be overestimated due to telomere rearrangements in these cells (Figure 3C). To gain further insight into the mechanisms at play, we measured the expression of both Taz1 and Rap1. At the protein level, we find that Taz1 is increased in *ccq1Δ* survivors (whether derived in the absence or presence of the GFP tag) and is maintained in RHR *ccq1Δ* survivors (Figure 5D), suggesting that enhanced expression may support the elevated levels of Taz1-GFP at the telomere in both types of survivors. By contrast, Rap1 expression levels remain nearly constant in all conditions (Figure 5D), arguing that altered recruitment, not less expression, underlies the loss of Rap1 from the telomere.

**Figure 5.**
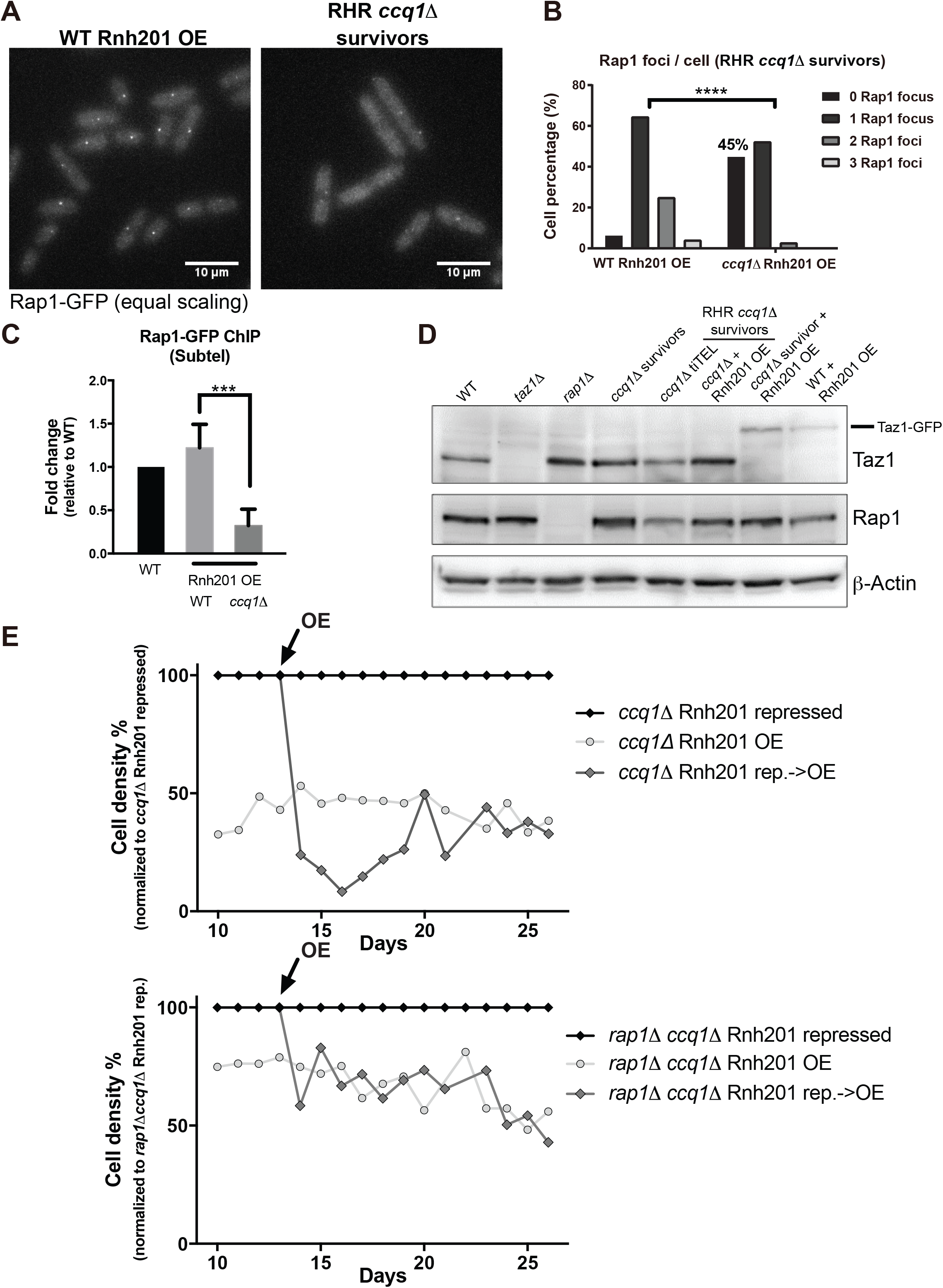
Loss of Rap1 promotes formation of RHR *ccq1Δ* survivors in the context of constitutive Rnh201 over-expression. (A) Fluorescence imaging of Rap1-GFP reveals decreased recruitment to the telomere upon long-term induction of Rnh201 in *ccq1Δ* survivor cells. Fluorescence micrographs of WT or *ccq1Δ* Rap1-GFP survivor cells after long-term Rnh201 over-expression (RHR *ccq1Δ* survivors). Maximum intensity projections obtained as described in the Methods. Scale bar = 10 μm (B) Quantification of the number of Rap1-GFP foci after long-term induction of Rnh201 in the WT and *ccq1Δ* backgrounds. At least 200 cells were analyzed. p < 0.0001 by K-S test. (C) The loss of Rap1 association with telomeres is also observed upon Rnh201 over-expression by chromatin immunoprecipitation-qPCR. Normalized to WT cells and plotted as the mean with SD from three biological replicates. *** p=0.0002 by Student’s t-test. (D) Taz1 levels increase in the absence of Rap1 and in all varieties of *ccq1Δ* survivors while Rap1 levels are unaffected. Whole cell extracts were separated on SDS-PAGE, blotted and probed with antibodies to Taz1, Rap1, or β-actin (as a loading control). (E) Deletion of Rap1 prevents the growth crisis upon Rnh201 over-expression in *ccq1Δ* survivors. Serial culturing of either *ccq1Δ* (top) or *rap1Δccq1Δ* (bottom) nmt41-Rnh201 cells under constitutive repression (black), over-expression (light gray circles), or induced induction of over-expression (OE at day 13, black arrow; dark gray diamonds). Cell density is normalized to the Rnh201-repressed condition.

### The growth crisis induced by RNA-DNA hybrid degradation can be ameliorated by deletion of Rap1

The loss of Rap1 from the telomere of RHR *ccq1Δ* survivors suggests that Rap1 may negatively influence TERRA-independent telomere maintenance by recombination. To test this hypothesis, we compared the growth response of *ccq1Δ* survivors to critical Rnh201 over-expression in the presence and absence of Rap1 (Fig. 5E). We carried out serial culturing of *ccq1Δ* nmt41-Rnh201 and *rap1Δccq1Δ* nmt41-Rnh201 cells under three conditions: constitutive Rnh201 repression (black, in the presence of thiamine), constitutive Rnh201 over-expression (OE; light grey circles) and switching from repression (13 days) to over-expression (arrow, dark grey diamonds). Consistent with Figure 3A,B, critical induction of Rnh201 in *ccq1Δ* cells at day 13 leads to a growth crisis (days 14-19), followed by recovery at day 20 to achieve a growth rate similar to cells cultured under conditions of constitutive Rnh201 over-expression. By contrast, critical Rnh201 over-expression in cells lacking Rap1 at day 13 led to a growth rate identical to cells under constitutive Rnh201 overexpression essentially immediately, without a growth crisis (Figure 5E and Figure S4A). We note that *ccq1* and *rap1* show a mild synthetic growth defect under conditions of Rnh201 repression (Figure S4B); to account for this, the plots in Fig. 5D-E represent the relative effect of RNA-DNA hybrid degradation. Raw growth plots reveal the same qualitative trend (Figure S4B). It has long been established that deletion of Rap1 leads to telomere elongation in telomerase-positive fission yeast cells {Kanoh, 2001 #404}, which is tied to enhanced recruitment of telomerase to the telomere {Dehe, 2012 #402;Moser, 2011 #360}. We therefore addressed the possibility that loss of Rap1 is able to compensate for defective telomerase recruitment in the absence of Ccq1, thereby supporting the growth of RHR *ccq1Δ* survivors through a telomerase-dependent mechanism. However, we find that deletion of the essential telomerase component Est1 has no effect on the growth of RHR *ccq1Δ* survivors (Figure S4C), consistent with previous work demonstrating that increased telomere length and telomerase recruitment in the absence of Rap1 is Ccq1-dependent {Moser, 2011 #360;Tomita, 2008 #1}. Thus, Rap1 appears to attenuate productive telomere recombination under conditions where RNA-DNA hybrids are degraded. How RHR *ccq1Δ* survivors displace Rap1 from the telomere despite its continued expression (Figure 5D) and the persistence of telomeric Taz1 (Figure 4), will require further study.

## DISCUSSION

Here we find that loss of the telomerase recruitment factor Ccq1 leads to progressive telomere shortening despite early loading of Rad52 onto telomeres, suggesting that additional factors are required for productive telomere lengthening by recombination.

We provide evidence that the spontaneous increase of telR-loop-engaged TERRA is a critical factor in the transition to productive telomere maintenance by recombination. In pre-senescent cells, increased telomere-associated TERRA is detrimental and telomere recombination is inefficient. By contrast, during survivor emergence, TERRA levels increase spontaneously (and/or are selected for during competition in liquid culture), form telR-loops, and promotes efficient telomere maintenance. Long-term degradation of telR-loops compromises the growth of *ccq1Δ* survivors, ultimately leading to a new survivor type with altered biochemical and cytological characteristics including loss of telomeric Rap1. Because deletion of Rap1 is sufficient to avoid the growth crisis induced upon RNA-DNA hybrid degradation in *ccq1Δ* survivors, we propose that TERRA-independent recombination pathways might be engaged to maintain telomeres by recombination.

### A switch from telomerase to recombination

As we place increased accumulation of telomere-associated TERRA as a key step in the transition to productive recombination, we favor a model in which changes to chromatin state within the subtelomere occur upon telomere shortening, supporting derepression of TERRA {Rippe, 2015 #443;Moravec, 2016 #423}. Indeed, induction of TERRA has been suggested to occur as part of the more global loss of the telomere position effect upon telomere shortening {Arnoult, 2012 #459;Mandell, 2005 #49;Cusanelli, 2013 #364}. Further, depletion of the histone chaperone ASF1 can induce a switch to the ALT pathway, suggesting that changes in the chromatin landscape are sufficient to activate HDR-dependent telomere maintenance {O’Sullivan, 2014 #355}. Ccq1 also contributes to recruitment of the SHREC complex (Sugiyama *et al.,* 2007) and the CLRC complex {Armstrong, 2018 #408;Wang, 2016 #407} to telomeres to promote heterochromatization. Disrupting the phosphorylation of Ccq1 by the DNA damage checkpoint kinases ATM and ATR, which enhances its interaction with Est1, abrogates telomerase recruitment without adversely affecting recruitment of SHREC {Moser, 2011 #360}; in this context survivors undergo chromosome circularization {Moser, 2011 #360}, suggesting that derepression of the subtelomere is critical for the maintenance of linear chromosomes by recombination. Combined with the observation that TERRA levels begin to increase after germination in cells lacking Ccq1 (Figure 2B), we propose that the combination of a more transcriptionally permissive chromatin state combined with shortened telomeres {Graf, 2017 #413} together drives high levels of TERRA, consistent with the ability of this genetic background to maintain linear chromosomes via recombination {Miyoshi, 2008 #42;Tomita, 2008 #1}. The slow emergence of survivors over many generations (and our failure to circumvent the growth crisis by TERRA expression from the tiTEL; Figure 2E,F) could reflect transmissibility of heterochromatic marks at the subtelomere and/or the requirement for additional changes, for example to checkpoint signaling, that remain to be uncovered.

### Support of recombination by telomere-associated TERRA

When telomere-associated TERRA is degraded, post-senescent *ccq1Δ* survivors reenter a growth crisis (Figure 3A,B). This finding is in line with the observation that TERRA is necessary for recombination at telomeres in budding yeast {Balk, 2013 #3}. Further, elevated levels of TERRA are characteristic of human ALT cell lines established from tumors or immortalized *in vitro* {Arora, 2014 #448;Lovejoy, 2012 #367}. TERRA could stabilize D-loop structures to facilitate recombination between telomeres {Balk, 2014 #357;Balk, 2013 #3}. Instead or in addition, TERRA transcription could promote an increase in telomere mobility to support telomere-telomere interactions necessary for recombination {Arora, 2012 #363}. Alternatively, it is possible that telR-loops driven by TERRA production promote break-induced replication, an HDR-based mechanism recently suggested to facilitate telomere maintenance in ALT cells {Dilley, 2016 #460}. While discriminating between these explicit molecular mechanisms will require further study, our data clearly support a model in which spontaneous derepression of TERRA plays a critical role in emergence of survivors in the absence of Ccq1.

### Distinct roles of TERRA in the presence or absence of telomerase

The effect that TERRA has on telomere length regulation varies, in part depending on the presence or absence of telomerase {Pfeiffer, 2012 #450;Redon, 2010 #461;Schoeftner, 2008 #440;Redon, 2013 #462;Balk, 2013 #3;Cusanelli, 2013 #364;Azzalin, 2008 #463}. In addition, our recent work revealed that TERRA molecules in fission yeast can be further grouped into at least two subsets: one is poly(A)+ TERRA, which loses all telomeric tracts and is soluble in the nucleoplasm; the other is poly(A)-TERRA, which has G-rich telomeric sequences, and is bound at telomeric DNA {Moravec, 2016 #423}. While poly(A)+ TERRA stimulates telomerase recruitment in telomerase-positive cells {Moravec, 2016 #423}, here we suggest that poly(A)-TERRA (or G-rich TERRA) promotes telomere recombination in cells with insufficient telomerase function. Therefore, these two subsets of TERRA molecules maintain telomeres through different mechanisms according to the cellular context. Expression of TERRA from a tiTEL cannot prevent *ccq1Δ* cell senescence, and in fact precipitates a more rapid growth crisis in pre-senescent cells (Figure 2E,F). This observation is consistent with the idea that increased TERRA production can be deleterious to telomere maintenance in some contexts {Balk, 2013 #3}. We note, however, that the tiTEL predominantly produces poly(A)+ TERRA and that nascent TERRA is limited to a single telomere; thus the tiTEL system may model the role for TERRA in telomerase-dependent {Moravec, 2016 #423} but not recombination-dependent telomere maintenance.

### TERRA dependent and independent recombination-based telomere maintenance

The “addiction” of *ccq1Δ* survivors to telomere-associated TERRA (suggested by the ability of Rnh201 over-expression to induce a second growth crisis) can be overcome, as evidenced by the formation of RNase H2-resistant *ccq1Δ* survivors. This mechanism is necessarily distinct from so-called HAATI survivors, which replace the telomeric repeats with “generic” heterochromatin, as this pathway requires the presence of Ccq1 {Jain, 2010 #31}. What changes might support a productive, TERRA-independent telomere recombination mechanism that can maintain telomeres in the absence of Ccq1? Our data suggest that these RHR *ccq1Δ* survivors display altered recruitment of telomere binding proteins, including loss of Rap1 (but not Taz1) from the telomere (Figures 4,5). This result is somewhat surprising, as Taz1 binds directly to Rap1 and promotes its recruitment to the telomere in fission yeast, thereby bridging the doublestranded telomeric repeat region to Poz1 and the shelterin components that associate with the single-stranded telomeric overhang {Pan, 2015 #464;Chikashige, 2001 #403;Kanoh, 2001 #404}. Moreover, loss of Taz1 and Rap1 phenocopy one another with respect to increased telomere length and de-repression of TERRA {Greenwood, 2012 #445;Bah, 2012 #366;Miller, 2005 #458}, although the mechanisms at play may be distinct {Dehe, 2012 #402;Miller, 2005 #458}. We note, however, that recent evidence supports allosteric regulation of Rap1 binding to Tpz1-Poz1 to regulate telomere length {Kim, 2017 #475}, suggesting that the Rap1-Taz1 interface may also be subject to modulation. Further studies will be required to test if changes in the composition and/or conformation of shelterin are tied to the mechanisms allowing for telomere maintenance in the absence of telR-loops.

Importantly, loss of Rap1 is not simply a consequence of the mechanisms that support telomere maintenance in RHR *ccq1Δ* survivors, because deletion of Rap1 ameliorates the growth crisis apparent upon degradation of RNA-DNA hybrids in *ccq1Δ* survivors (Figure 5E). How Rap1 eviction from the telomeres promotes the maintenance of linear chromosomes in this context is not yet clear, although we have ruled out the participation of telomerase (Figure S4C). Given that Rap1 is a known repressor of TERRA {Bah, 2012 #366}, one possibility is that there is a further stimulation of TERRA production beyond that seen in survivors that is able to restore telomere-engaged TERRA. Alternatively, decreased telomeric Rap1 could weaken the association of telomeres with the nuclear envelope, which has been suggested to repress telomere recombination in budding yeast {Schober, 2009 #465}. Ultimately, testing how changes in shelterin composition may act independently of TERRA to promote telomere maintenance by ALT will require further study.

## Acknowledgements

We are indebted to the Yeast Genomic Resource Center (YGRC) at Osaka University for providing access to strains, as well as the many researchers who have deposited strains at this resource. We also thank J. Kanoh and J. P. Cooper for sharing antibodies.

