## Supplemental Table 1, Supplemental Figs. 1-4 for "RNA-DNA hybrids support recombination-based telomere maintenance in fission yeast"

Supplemental Materials: Table S1 and Figures S1-S4 with Legends

**Supplemental Figure Legends:**

**Figure S1.** (A) Biological replicate of serial culturing of WT, *ccq1Δ*, and *ccq1Δrad55Δ* cells as in Figure 1A. (B) Southern blot of EcoRI digested genomic DNA from WT cells or *ccq1Δ* cells at 1, 18 or 30+ days in liquid culture to give rise to survivors (S), hybridized with a DIG-labeled telomere probe derived from a cloned telomere and visualized using a biotinylated anti-DIG antibody followed by streptavidin-HRP and chemilluminescence imaging. The contrast was increased to visualize telomere signal in the bottom panel. The *ccq1Δ* survivors also have a higher band of ~ 8 kb, likely due to the partial loss of the distal EcoRI cutting site at telomere ends, and these telomeres are therefore restricted at the second, upstream EcoRI site instead. (C) TERRA is induced in *trt1Δ* cells but to a lesser extent than seen in *ccq1Δ* cells. Total TERRA levels in *trt1Δ* cells after 20, 26 or over 200 generations in liquid culture (survivors) as determined by RT-PCR as in Figure 2A.

**Figure S2.** (A) Validation of Rnh201 over-expression upon removal of thiamine. Anti-HA Western blot of WT Taz1-GFP or *ccq1*Δ Taz1-GFP strains with nmt41-3xHA integrated before the start codon of the Rnh201 gene. Samples were prepared with and without thiamine for 24 hours, as indicated. The signal from 3xHA-Rnh201 is indicated with an arrow. The asterisk indicates a non-specific band that is also visible in WT cells without any HA integration. (B) nmt41-driven Rnh201 is transcribed at levels higher than WT cells in the presence of thiamine, with a further increase in Rnh201 expression upon thiamine removal. The mRNA level of Rnh201 was detected by quantitative RT-PCR in WT cells or WT cells with the inducible Rnh201 construct (*nmt41-3HA-Rnh201*) grown in minimal medium with or without thiamine. (Rnh201-RT: 5’-CTCTGTAACATTTATTCCTGATTCATAAAC-3’; Rnh201-For: 5’-TTGACCAACTATGGGTTTGC-3’; Rnh201-Rev: 5’- CTGCAGCTAATTCTCTTGCG-3’). (C) Rnh201 over-expression reduces telomeric RNA/DNA hybrids in WT and *ccq1*Δ survivors. Cells with an inducible Rnh201 construct (*nmt41-3HA-Rnh201*) were collected under different conditions: grown in EMM5S medium plus 5 μg/ml thiamine (No induction), grown in EMM5S medium for 16 hours (Rnh201 OE 16h), and grown in EMM5S medium for 15 days with serial dilution each day (Rnh201 OE 15d). (D) Induction of 3xHA-Rnh201 in *ccq1*Δ survivors for 24 hours grow in EMM leads to loading of Rad52-mCherry at the telomere (labeled by Taz1-GFP; arrowheads) and anaphase bridges (arrow). Fluorescence micrographs of strain MKSP2540. Scale bar = 5 μm. (E) 3xHA-Rnh201 over-expression is maintained in RHR *ccq1Δ* survivors. Western blots were carried out as described in (A).

**Figure S3.** (A-B) The appearance of Taz1-GFP in *ccq1*Δ survivors and RHR *ccq1*Δ survivors is the same if Taz1-GFP is integrated after survivor formation. Maximum intensity projections of fluorescence micrographs of *ccq1*Δ survivors and RHR *ccq1*Δ survivors after integration of Taz1-GFP obtained as described in the Methods. Scale bar = 5 μm (C) Deletion of Rap1 does not affect the appearance of Taz1-GFP. Maximum intensity projections of fluorescence micrographs obtained as described in the Methods. Scale bar = 1 μm (D) and (E) Recruitment of Rap1-GFP to the telomere is not significantly altered in *ccq1*Δ survivors. (D) The intensity of focal Rap1-GFP signal is not statistically different between WT and *ccq1*Δ survivors. ns = not significant (E) There is only a subtle shift to fewer Rap1-GFP foci per cell in WT compared to *ccq1*Δ survivors. n>200 cells for each genotype. p = 0.03 by K-S test.

**Figure S4.** (A) Deletion of Rap1 prevents the growth crisis upon Rnh201 over-expression in *ccq1Δ* survivors. Additional biological replicate of the *rap1Δccq1Δ* nmt41-Rnh201 growth assay under constitutive repression (black), over-expression (light gray circles), or induced induction of over-expression (OE at day 13, black arrow; dark gray diamonds). (B) Raw cell density plots for the serial dilution assays in Figure 5E for *ccq1Δ* (top) or *rap1Δccq1Δ* (bottom) nmt41-Rnh201 cells under constitutive repression (black), over-expression (light gray circles), or induced induction of over-expression (OE at day 13, black arrow; dark gray diamonds). (C) Deletion of Est1 does not compromise growth of RHR *ccq1*Δ survivors. Serial 10-fold dilutions of RHR *ccq1*Δ survivors or RHR *ccq1*Δ survivors transformed to remove Est1 grown on YE5S media.

**Table S1: Fission yeast strains and strains used in their derivation in this study, as well as survivors generated by serial culturing.**

| Name (MKSP) and Fig. used | GENOTYPE | SOURCE |
|---|---|---|
| 110 **Fig. 5D** | *rap1::ura4+ leu1-32 h-* | YGRC FY14171 |
| 112 **Fig. 5D** | *taz1::ura4+ leu1-32 h-* | YGRC FY14161 |
| 200 | *ade6-M216 ura4-D18 leu1-32 h+* | {Nurse, 1976 #95} |
| 291 **Fig. 3D** | *ade6-M210 leu1-32 ura4-D18 trt1::LEU2 h90* | YGRC FY14084 |
| 310 | *ccq1::HygR leu1-32 ura4-D18 h-* | TM2370 {Miyoshi, 2008 #42} |
| 399 **Fig. 2E, Fig. S1B** | *ura4-D18 leu1-32 h+* ("WT") | this work |
| 581 **Fig. S3D,E** | *Rap1-GFP::NatR leu1-32 ura4-D18 h+* | this work |
| 863 **Fig S1A, 2C, 2D** | *ura4-D18/ura4-D18 leu1-32/ leu1-32 ade6-M210/ade6-M216 ccq1::HygR/CCQ1 rad55::NatR/RAD55 h+/h+* | this work |
| 891 **Fig. 2D, 3E** | *ura4-D18 leu1-32 ade6-M21? ccq1::HygR rad55::NatR h+* | this work, survivor |
| 1027 **Fig. 2E** | *ccq1:HygR leu1-32 ura4-D18 h+* | this work, derived from MKSP310 |
| 1158 | *ura4-D18 mcp5::ura4+ Taz1-GFP::KanR h90* | YGRC FY16842; ST242-1, Nojima lab (Osaka University) |
| 1165 **Fig. S1C** | *poz1+/poz1:HygR trt1+/trt1::KanR ade6+/ade6-M216 leu1-32/leu1-32 h+/h-* | TM2571 (Miyoshi *et al.*, 2008) |
| 1211 **Fig. 2A, Fig. S1A,B** | *ccq1::HygR leu1-32 ura4-D18 h+* | this work, survivor |
| 1213 **Fig. 4** | *ura4-D18 Taz1-GFP::KanR leu? ade? h?* | this work, derived from MKSP1158 |
| 1214 **Fig. 1B,D; Fig. 2A,B, Fig. 4, Fig. 5D** | *ccq1::HygR leu? ura4-D18 Taz1-GFP::KanR h+* | this work; survivors |
| 1223 **Fig. 1B,D** | *ura4-D18 Taz1-GFP::KanR leu? ade? Rad52-mCherry:natMX6 h+* | this work |
| 1262 **Fig. S3A** | *ccq1::HygR leu1-32 ura4-D18 ade6-M21? Taz1-GFP::KanR h? (Taz1-GFP integrated after survivor formation)* | this work, survivor |
| 1579 **Fig. 3, 5D, S2** | *ura4-D18 Taz1-GFP::KanR leu? ade? NatR::nmt41-3HA-Rnh201 h?* | this work |
| 1580 **Fig. 3, S2** | *ccq1::HygR leu? ura4-D18 Taz1-GFP::KanR NatR::nmt41-3HA-Rnh201 h+* | this work, survivor |

| | | |
|---|---|---|
| 1672 **Fig. 2E** | *NatR::P3nmt1-Tel-I-R* | CAF544 (Moravec *et al.*, 2016) |
| 1709 **Fig. 4, 5D** | *ccq1::hygR leu? ura4-D18 Taz1::GFP::KanR NatR::nmt41-3HA-Rnh201 h?* | this work, *ccq1∆* survivor with Rnh201 OE starting from spore with Taz1-GFP expression |
| 1886 **Fig. 5** | *leu1-32 ura4-D18 Rap1-GFP-Kan nmt41-3HA-Rnh201::NatR h?* | this work |
| 1887 **Fig. S3D,E** | *leu1-32 ura4-D18 Rap1-GFP ccq1::HygR h+* | Rap1-GFP transformed into *ccq1∆* survivor |
| 1902 **Fig. 2E, 2F, 5D** | *ccq1::hygR NatR::P3nmt1-Tel-I-R h?* | this work |
| 1934 **Fig. 5** | *ccq1::hygR leu1-32 ura4-D18 ade6-M21? Rap1::GFP-KanR nmt41-3HA-Rnh201::NatR h?* | this work ("RHR survivor" with Rap1-GFP) |
| 1988 **Fig. S3C** | *rap1::ura4 Taz1-GFP::KanR ura4-D18 leu1-32 h?* | this work |
| 1992 | *ccq1::hygR leu? ura4-D18 NatR:nmt41-HA-Rnh201 h?* | this work, *ccq1∆* survivor with Rnh201 OE starting from spore |
| 2027 **Fig. 2F** | *NatR::P3nmt1-Tel-I-R ccq1::hygR Taz1::GFP::KanR Rad52-mCherry:KanR ura4-D19 leu1-32 h?* | this work, derived from MKSP2059/2060 |
| 2059 | *Rad52-mCherry:KanR ura4-D18 leu1-32 his3-D1 h-* | this work |
| 2060 | *NatR::P3nmt1-Tel-I-R ccq1::hygR Taz1::GFP::KanR h+* | this work |
| 2497 **Fig. S3B** | *ccq1::hygR leu? ura4-D18 Taz1::GFP::KanR NatR:nmt41-HA-Rnh201 h? Taz1-GFP integration post-selection of survivors* | this work, generated from MKSP1992 |
| 2540 **Fig. 3C, Fig. S2D** | *ccq1::hygR leu? ura4-D18 Taz1::GFP::KanR Rad52-mCherry::KanR NatR:nmt41-HA-Rnh201 h+* | this work |
| 2762 **Fig. 5E, S4A,B** | *RAP1/rap1::ura4+ leu1-32/leu1-32 ura4-D18/ura4-D18 CCQ1/ccq1::hygR RNH201/nmt41-3HA-Rnh201::NatR h+/h+* | this work |
| 2820 **Fig. S4C** | *ccq1::hygR leu1-32 ura4-D18 est1::KanR nmt41-3HA-Rnh201::NatR h?* | this work, derived from MKSP1992 |
| 2983 **Fig. 1A,C** | *ura4-D18/ura4-D18 leu1-32/ leu1-32 ade6-M210/ade6-M216 ccq1::HygR/CCQ1 rad55::NatR/RAD55 h+/h+ Rad52-mCherry::KanR* | this work |

**References:**


Moravec, M., Wischnewski, H., Bah, A., Hu, Y., Liu, N., Lafranchi, L., King, M. C., Azzalin, C. M. (2016). TERRA promotes telomerase-mediated telomere elongation in *Schizosaccharomyces pombe*. EMBO Rep *17*, 999-1012.

Miyoshi, T., Kanoh, J., Saito, M., and Ishikawa, F. (2008). Fission yeast Pot1-Tpp1 protects telomeres and regulates telomere length. Science *320*, 1341-1344.

Nurse, P., Thuriaux, P., and Nasmyth, K. (1976). Genetic control of the cell division cycle in the fission yeast Schizosaccharomyces pombe. Molecular & general genetics : MGG *146*, 167-178.

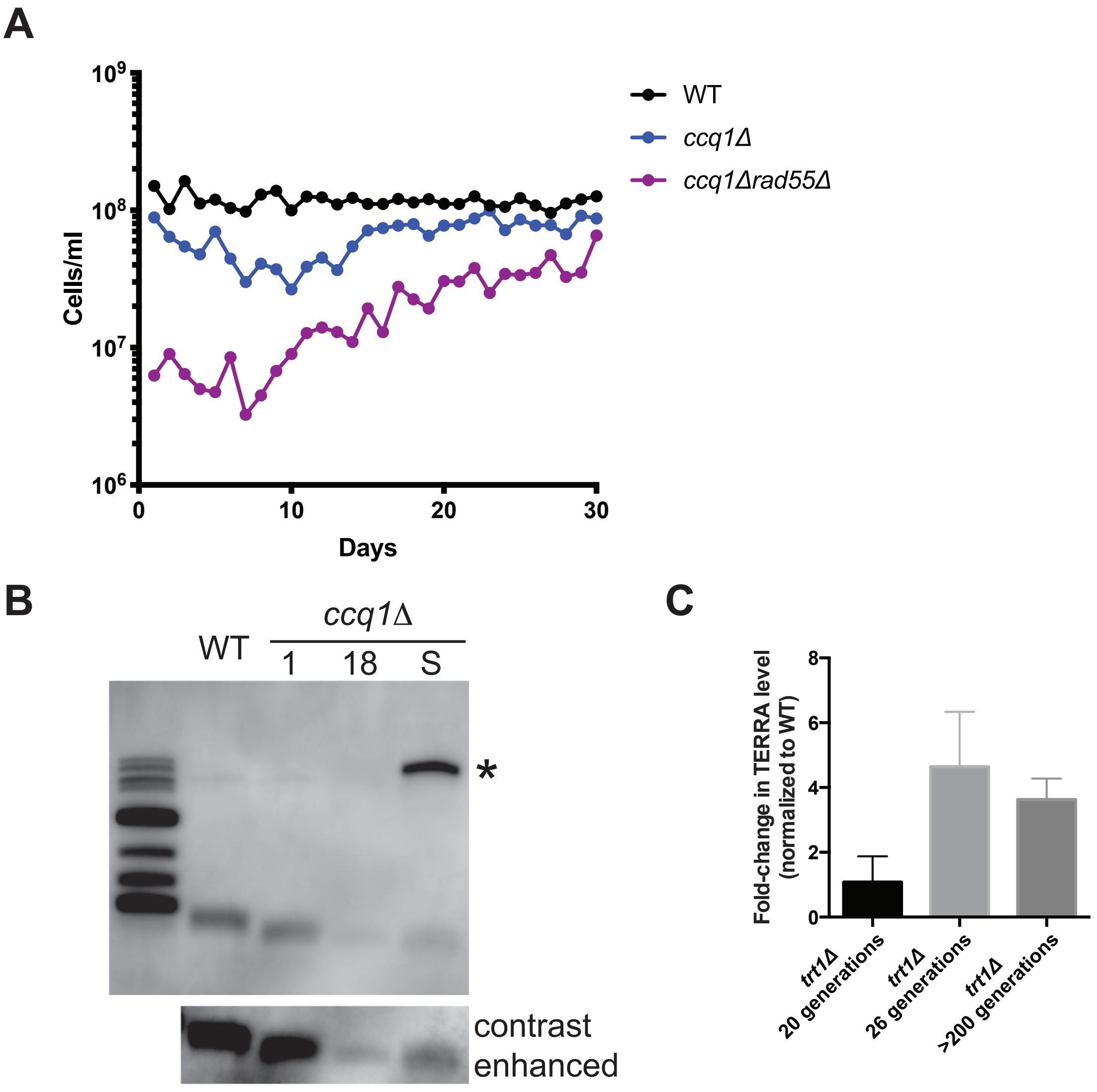


Hu et al., Figure S1

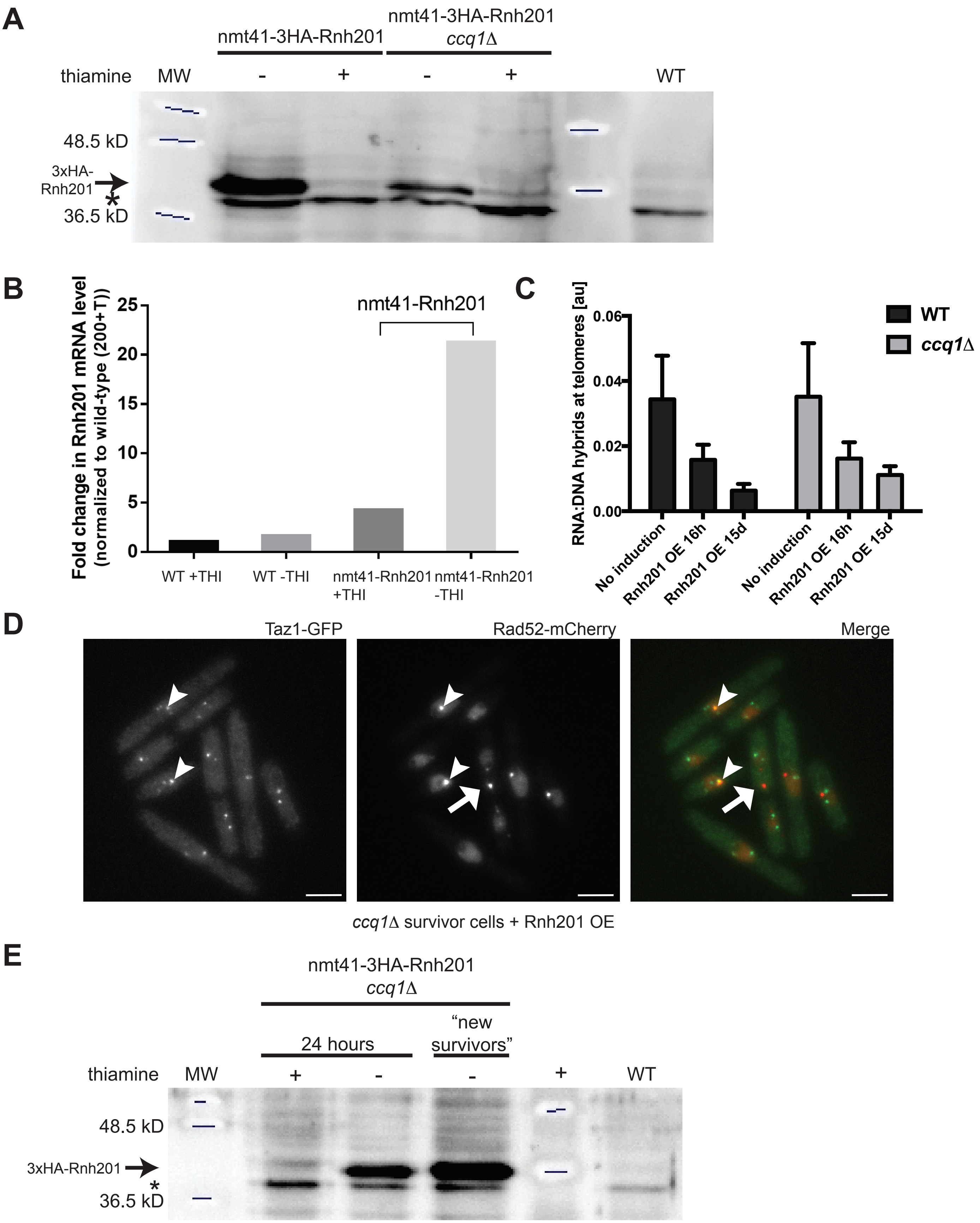


Hu et al., Figure S2

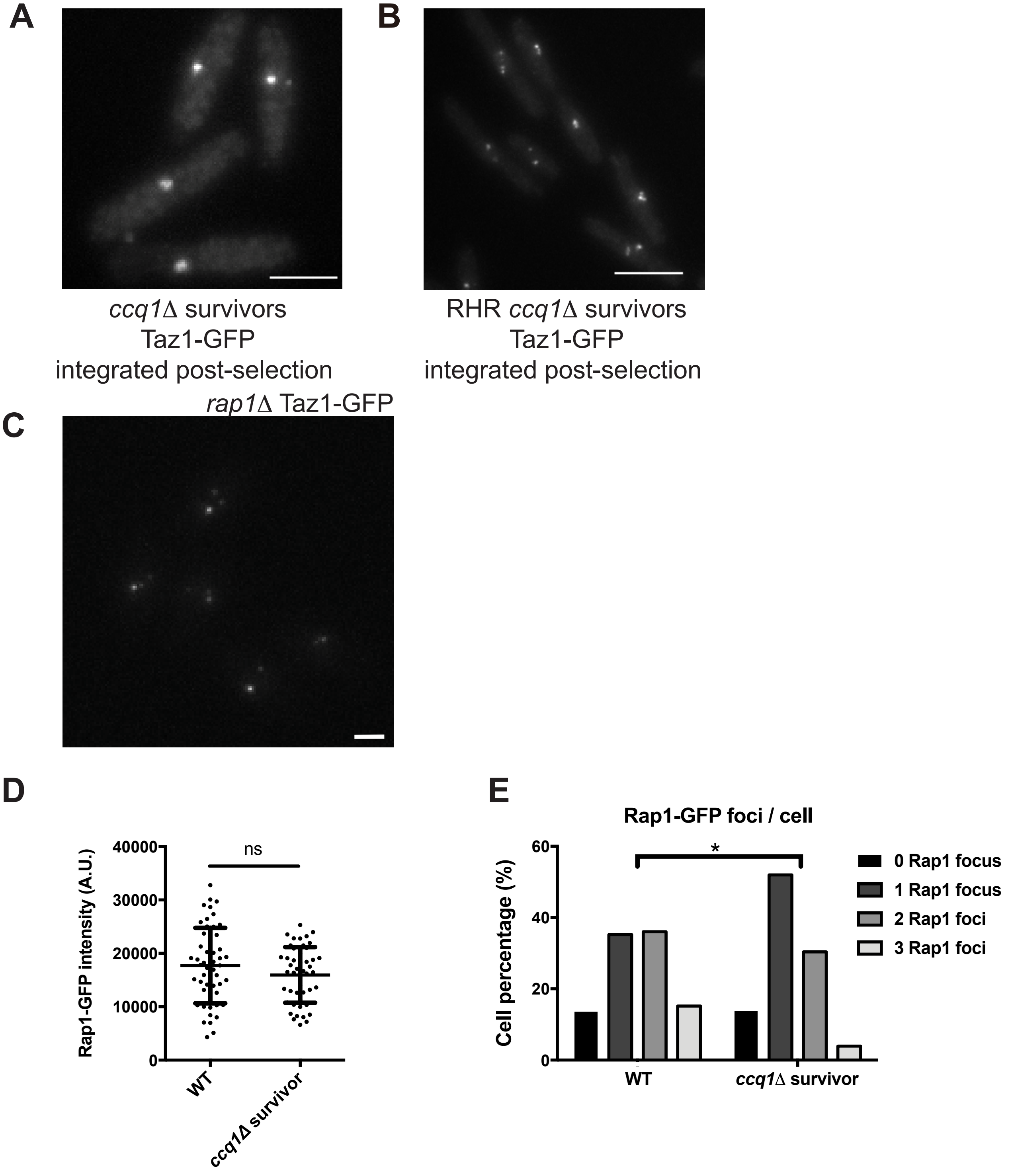


Hu et al., Figure S3

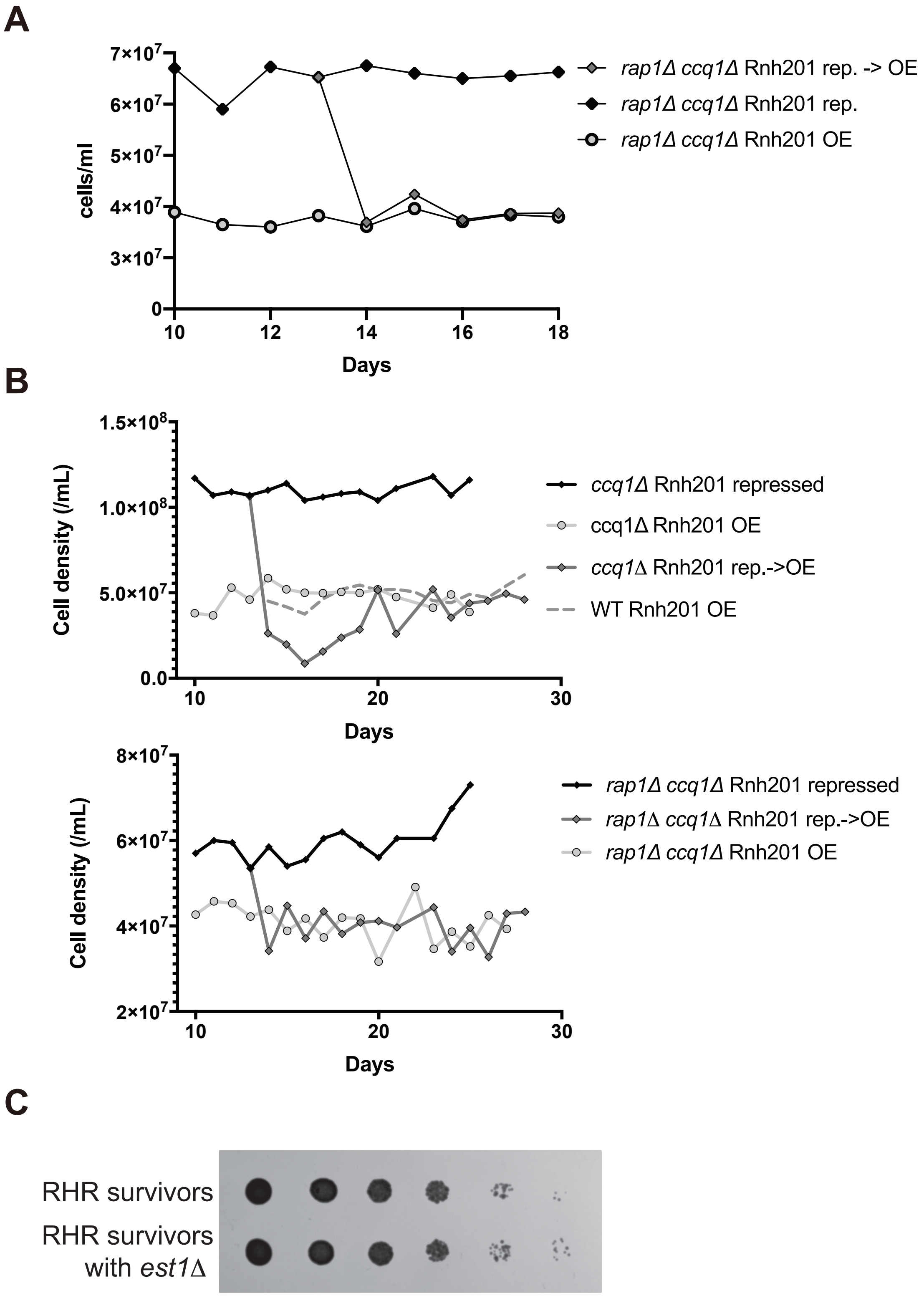


Hu et al., Figure S4
